# N-Acylethanolamine is a Source of Ethanolamine for Phosphatidylethanolamine Synthesis in *Trypanosoma brucei*

**DOI:** 10.1101/2023.07.20.549923

**Authors:** Aurelio Jenni, Peter Bütikofer

## Abstract

The Kennedy pathway is essential for survival of *Trypanosoma brucei* procyclic and bloodstream form parasites in culture and provides the bulk of phosphatidylethanolamine and phosphatidylcholine for membrane biogenesis. The CDP-ethanolamine branch of the Kennedy pathway depends on a steady supply of ethanolamine to be functional. We now show that degradation of N-acylethanolamines, mediated at least in part by *T. brucei* fatty acid amide hydrolase (TbFAAH), represents an additional reaction to provide ethanolamine for phosphatidylethanolamine synthesis. Although TbFAAH is not essential for growth of *T. brucei* procyclic forms in normal culture medium, it may be essential for ethanolamine production under conditions of limited availability of free ethanolamine in the environment, or in the insect vector or mammalian host.

## Introduction

The glycerophospholipid phosphatidylethanolamine (PE) is not only a major building block of eukaryotic membranes (1), but has also other important roles in a variety of physiological processes, including protein folding (2), oxidative phosphorylation (3) and autophagy (4). In addition, PE serves as the ethanolamine donor for the glycosylphosphatidylinositol (GPI) anchor biosynthesis pathway (5, 6) and is the precursor of the ethanolamine phosphoglycerol modification of eukaryotic elongation factor 1A (7).

In the protozoan parasite, *Trypanosoma brucei*, PE represents the second most abundant glycerophospholipid class (8, 9). Its synthesis occurs via the CDP-ethanolamine branch of the Kennedy pathway (10), which represents the only pathway for *de novo* synthesis of PE in *T. brucei* (11) and, thus, is essential for parasite survival in culture (11–13). First, ethanolamine is phosphorylated followed by activation of ethanolamine-phosphate to CDP-ethanolamine. Subsequently, CDP-ethanolamine is conjugated with diacylglycerol or alk-1-enyl-acylglycerol to form diacyl- or ether-type PE molecular species, respectively. Acylation of *lyso*-PE taken up from the environment (14, 15) is another pathway to generate PE, its contribution to the total pool of PE, however, is unclear (7, 13). In addition, PE can also be generated by head group exchange with (16) or decarboxylation of phosphatidylserine (PS) (11, 13). The former pathway is fully reversible and provides total PS in *T. brucei* (17). Thus, PE produced from PS by either pathway does not contribute to *de novo* PE synthesis.

Ethanolamine and ethanolamine-phosphate are key precursors for PE formation via the Kennedy pathway, implying that a steady supply is essential for PE synthesis. *T. brucei* parasites have developed several strategies to obtain ethanolamine or ethanolamine-phosphate. Labeling experiments have demonstrated that ethanolamine is rapidly taken up from the environment by *T. brucei* parasites and incorporated into PE via the Kennedy pathway (18). This route provides the majority of ethanolamine for PE production. In addition, ethanolamine-phosphate can be generated via degradation of sphingosine-1-phosphate by sphingosine-1-phosphate lyase (19) (20). Furthermore, ethanolamine can be generated by decarboxylation of serine, however, while this pathway is active in other parasites, such as *Plasmodium falciparum* (21), it seems absent in *T. brucei*. Finally, it has been proposed (22), but not experimentally proven, that ethanolamine may also be generated via hydrolysis of N-acylethanolamines (NAEs) by fatty acid amide hydrolases (FAAHs) (23) or N-acylethanolamine-hydrolyzing acid amidases (NAAAs) (24, 25), reactions releasing free fatty acids and ethanolamine.

NAEs are a class of lipids with signaling properties. The conserved structure consists of a fatty acid linked to the nitrogen atom of ethanolamine. The fatty acid moiety is variable and defines the physiological properties of NAEs. Examples include anandamide (arachidonoylethanolamide, AEA), a well-studied endocannabinoid and endogenous agonist of the cannabinoid receptors (26), and palmitoylethanolamide (PEA), an anti-inflammatory NAE (27). Both AEA and PEA are present in significant quantities in human blood plasma (28) and – perhaps more important when working with cell cultures – in fetal bovine serum (29). In mammals, the physiological activity of NAEs is terminated through hydrolysis by FAAH or NAAA enzymes.

In this study, we characterized a putative *T. brucei* FAAH-2 homologue (TbFAAH) and investigated the parasite’s ability to generate ethanolamine via NAE hydrolysis. Our results show that *T. brucei* take up and hydrolyze AEA and use the generated ethanolamine for PE synthesis via the Kennedy pathway. In *T. brucei* TbFAAH knockout parasites, NAE hydrolysis was decreased, but not absent, and parasite growth in culture was not affected.

## Materials and Methods

Unless otherwise stated, reagents were purchased from MilliporeSigma (Burlington, MA, USA) or Merck KGaA (Darmstadt, Germany). Restriction enzymes were from ThermoFisher Scientific (Waltham, MA, USA). [^3^H]-labeled anandamide ([^3^H]-AEA, with the label being in the ethanolamine moiety; 60 Ci/mmol) was from American Radiolabeled Chemicals Inc. (St. Louis, MO, USA) and PCR reagents and restriction enzymes from Promega Corporation (Madison, WI, USA). Acrylamide mix was from National Diagnostics (Atlanta, GA, USA).

### Target identification

Possible NAE-degrading enzymes were identified using the pBLAST tool of the National Center for Biotechnology Information (NCBI). pBlast searches with the amino acid sequences of human FAAH-1, FAAH-2 and NAAA were performed against the deduced protein sequences of *T. brucei*.

### Trypanosome Cultures

*T. brucei* SmOx P9 pTB011 procyclic forms (30) (henceforward SmOx P9) were maintained at 27 °C in SDM79 containing 10% (v/v) heat-inactivated fetal bovine serum, 160 μM hemin, 90 μM folic acid, 2 μg/ml puromycin and 5 μg/ml blasticidin. Knock-out (KO) parasites were grown under the same conditions with 25 µg/ml hygromycin added as selection antibiotic. *T. brucei* 29-13 procyclic forms (31) (henceforward 29-13) were maintained at 27 °C in SDM79 containing 10% (v/v) heat-inactivated fetal bovine serum, 160 μM hemin, 90 μM folic acid, 25 μg/ml hygromycin and 1 μg/ml G418. Cells containing a tetracycline-inducible, cMyc-tagged copy of TbFAAH were grown under the same condition with 2 μg/ml puromycin added as a selection antibiotic. Protein expression was induced by addition of 0.1 μg/ml tetracycline. *T. brucei* New York Single Marker bloodstream forms (31) (henceforward NYSM) were maintained at 37 °C in HMI-9 containing 10% (v/v) heat-inactivated fetal bovine serum, 160 μM hemin, 90 μM folic acid and 2.5 μg/ml G418. Cells containing a tetracycline-inducible, HA-tagged copy of TbFAAH were grown under the same condition with 2.5 μg/ml puromycin added as a selection antibiotic. Protein expression was induced by addition of 0.1 μg/ml tetracycline.

### Generation of *T. brucei* TbFAAH-KO Parasites

Procyclic form TbFAAH-KO parasites were generated using clustered regularly interspaced short palindromic repeats (CRISPR) / CRISPR-associated protein 9 (Cas9) technique as described before (30). Briefly, a resistance gene cassette was generated by PCR using primers 1 and 2 (Table S1) and template plasmid pPOTv6 (32) containing the resistance gene for hygromycin. The cassette was flanked with homology sequences of 30 nt to replace both alleles of the target gene via homologous recombination. Two single-guide RNA templates containing a T7 polymerase promoter, a Cas9 binding site and a 20 nt targeting sequence were generated by PCR using primer pairs 3/5 and 4/5 (Table S1), respectively. All PCRs were performed using the Expand High Fidelity PCR System (Roche Diagnostics GmbH, Mannheim, Germany) and primers were designed using the online tool at www.leishgedit.net. All PCR products were pooled and purified using the Wizard® SV Gel and PCR Clean-Up System (Promega). DNA (10 μg) was transfected into SmOx P9 cells using a 4D nucleofector system (Lonza Group AG, Basel, Switzerland) with program FI-115. After 24 h, selection antibiotics were added, and the cultures were diluted 1:25 and distributed into 24-well plates. TbFAAH gene knock-out was verified by PCR using extracted gDNA from knock-out clones and primer pair 6/7 (Table S1).

### Generation of *T. brucei* Parasites Overexpressing TbFAAH-HA

*T. brucei* 29-13 and NYSM parasites containing a tetracycline-inducible HA-tagged copy of TbFAAH were generated using *Not*I-linearized transfection constructs derived from plasmids pALC14 (a kind gift of André Schneider, University of Bern, Switzerland) for 29-13 and pMS (33) for NYSM.

### Immunofluorescence Microscopy

Approximately 2.5 × 10^6^ trypanosomes were harvested by centrifugation, washed with cold phosphate-buffered saline solution (PBS; 137 mM NaCl, 2.7 mM KCl, 10 mM Na_2_HPO_4_, 1.76 mM KH_2_PO_4_, pH 7.4), resuspended in a small volume of PBS, spread on a microscopy slide and left to adhere for 20 min. Subsequently, parasites were fixed using 4% (w/v) paraformaldehyde in PBS for 10 min. After three washes for 5 min with cold PBS, cells were permeabilized in 0.2% (w/v) Triton X-100 for 20 min. After three additional washes and incubation in blocking solution (2% (w/v) bovine serum albumin in PBS) for 30 min, the cells were incubated with primary antibody in blocking solution for 45 min at room temperature. Antibodies used were mouse α-HA.11 (Covance, Princeton, NJ, USA) and rabbit α-BiP (provided by James D. Bangs, University at Buffalo, Buffalo, NY, USA) at dilutions of 1:500 and 1:2500, respectively. After washing, the cells were incubated for 45 min with fluorophore-conjugated secondary antibody goat α-mouse AlexaFluor594 or goat α-rabbit AlexaFluor488 at 1:1000 dilutions in blocking solution. After washing, the cells were mounted with Vectashield DAPI (Vector Laboratories, Burlingame, CA, USA). Immunofluorescence image stacks were captured on a Leica SP2 using a 100× oil objective. Image stacks were 3D deconvolved with the Leica LAS AF Version 2.1.0 software (Leica Microsystems CMS GmbH, Heerbrugg, Switzerland).

### Thin-Layer Chromatography (TLC)

Samples were spotted onto Silica Gel 60 plates using glass capillaries. The plates were developed in chloroform/methanol/acetic acid/water (25:15:4:2; by vol.), dried and analyzed using a Raytest Rita* radioactivity TLC analyser (Berthold Technologies, Regensburg, Switzerland).

### Determination of Spontaneous [^3^H]-Anandamide Hydrolysis

[^3^H]-AEA (3.3 pmol) was added to 500 μl culture medium and incubated for 24 h at 27 °C (for SDM79) or 37 °C (for HMI-9). Subsequently, 1 ml of chloroform/methanol 1:1 (v/v) was added to each sample, and the solutions were vortexed. Phase separation was achieved by centrifugation at 17000 x g for 1 min. Aliquots of 100 μl of each phase were used for analysis by TLC.

### [^3^H]-PE Formation Using [^3^H]-Labeled Anandamide

Trypanosomes in logarithmic growth (4×10^7^ cells) were harvested by centrifugation and resuspended in 4 ml culture medium containing 1.5 µCi [^3^H]-AEA. After 4 h of incubation at 27 °C (for procyclic forms) or 37 °C (for bloodstream forms), parasites were centrifuged, washed with TBS and hypotonically lysed by the addition of 1 ml of water. Phospholipids were extracted according to Bligh and Dyer (35). Briefly, the lysate was transferred to a glass tube and 3.75 ml chloroform/methanol 1:2 (v/v) was added. After 30 min on ice, phase separation was induced by addition of 1.25 ml water and 1.25 ml chloroform. After vortexing thoroughly, the lower phase, containing the (phospho-) lipids, was collected, dried, and analyzed by TLC.

### FAAH Activity Assay

FAAH activity was measured as described elsewhere (36). Briefly, trypanosomes (10^8^ cells/sample) were harvested by centrifugation and resuspended in 500 µl FAAH activity buffer (10 mM Tris-HCl, 1 mM EDTA, 1 mg/ml bovine serum albumin, pH 8.0). Parasites were lysed by freezing at -20 °C for 24 h followed by homogenization through a G27 needle. AEA mix (2 µl of 99.5 nM AEA, containing 0.5 nM [^3^H]-AEA, in ethanol) was added and the samples were incubated for 15-120 min at 27 °C. The reaction was stopped by the addition of 1 ml chloroform/methanol 1:1 (v/v). After vortexing thoroughly, phases were separated by brief centrifugation. Aliquots of 600 µl of both phases were removed, dried and analyzed by liquid scintillation counting. Appearance of radioactivity in the aqueous phase was taken as indication of FAAH hydrolysis activity.

## Results and Discussion

### The *T. brucei* genome contains a homologue of human FAAH-2

pBLAST searches identified a predicted 595 amino acid protein with a calculated molecular mass of 65 kDa, designated TbFAAH and encoded by the gene Tb927.8.5080, with an identity of 38% on the amino acid level and an e-value of 10^−106^ against human FAAH-2. In analogy to human FAAH-2 (37), the predicted *T. brucei* protein contains an amidase signature sequence between amino acids 87 and 344 (38) and a single N-terminal transmembrane domain between amino acids 12 and 35. Based on TrypTag.org (39), TbFAAH localizes to the nuclear envelope.

### TbFAAH is not essential for growth of *T. brucei* procyclic forms

To study the importance of TbFAAH for survival of *T. brucei* procyclic forms in culture, the two alleles of Tb927.8.5080 were targeted for replacement by antibiotic resistance cassettes. During the first round of transfection, exchange of one allele with a hygromycin resistance cassette was attempted. Two hygromycin-resistant clones were selected and screened for allele replacement by PCR (Fig. 1). The first clone, designated TbFAAH-sKO, revealed the disappearance of one of the two Tb927.8.5080 alleles and the presence of the hygromycin resistance gene (Fig. 1), indicating that one allele has been replaced by a hygromycin resistance gene. Unexpectedly, the second hygromycin-resistant clone, designated TbFAAH-KO, showed the absence of both Tb927.8.5080 alleles and the presence of a single strong band representing the hygromycin resistance gene (Fig. 1), suggesting that both alleles of Tb927.8.5080 were exchanged simultaneously by hygromycin resistance genes during this first round of transfection.

**Figure 1:**
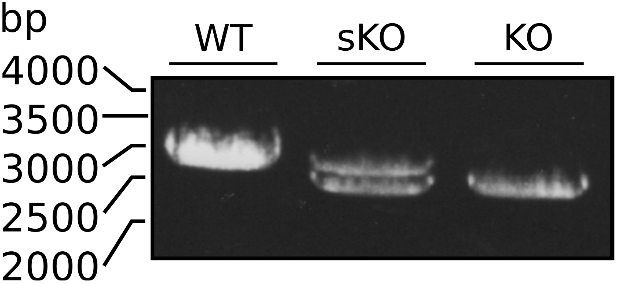
Analysis of TbFAAH-KO clones. gDNA was extracted from SmOx P9 (WT), TbFAAH-sKO and TbFAAH-KO cells. Amplification of the TbFAAH locus was done by PCR using primers 6 and 7, binding to the untranslated regions flanking the TbFAAH locus (Table S1).

The importance of TbFAAH for *T. brucei* growth was analyzed by determining parasite proliferation in culture. The results show that growth of SmOx P9 (parental cells) and TbFAAH-KO cells was similar, with cell doubling times of 9.4 and 9.8 h, respectively, indicating that deletion of the TbFAAH gene (Tb927.8.5080) had no major effect on growth of *T. brucei* procyclic forms in culture (Fig. 3).

**Figure 3:**
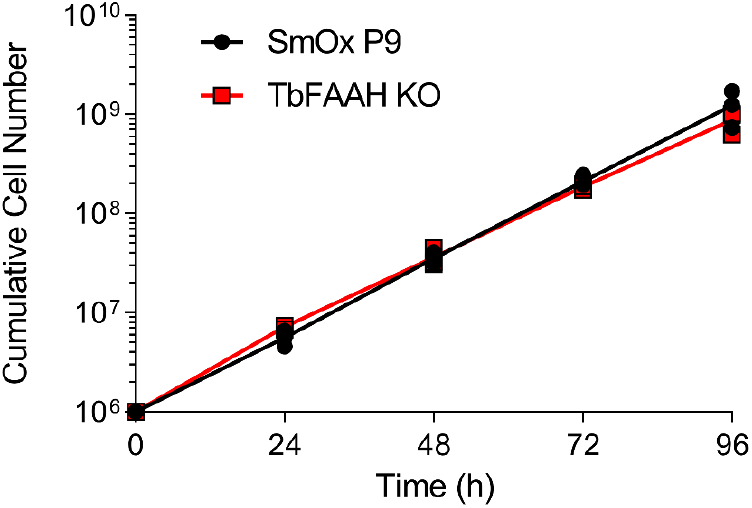
Growth of TbFAAH-KO parasites. T. brucei wild-type (SmOx P9) and TbFAAH-KO procyclic forms were cultured in standard medium and parasite densities were recorded every 24 h over 4 days. Data are from 3 independent experiments.

### HA-tagged TbFAAH localizes to the ER

To study the localization of TbFAAH, a tetracycline-inducible HA-tagged copy of the enzyme (TbFAAH-HA) was expressed in *T. brucei* procyclic and bloodstream forms. Analyses by SDS-PAGE and immunoblotting showed that TbFAAH-HA was expressed in both life cycle forms after induction by tetracycline with an apparent molecular mass of approximately 60 kDa (Fig. 4).

**Figure 4:**
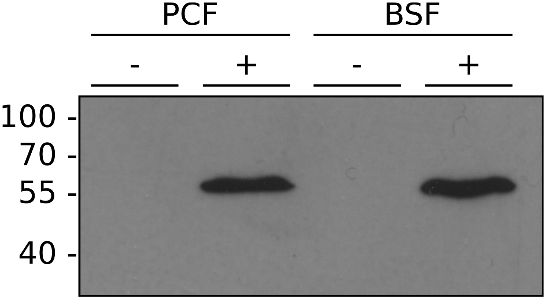
Expression of TbFAAH-HA. T. brucei procyclic (PCF) and bloodstream (BSF) forms were cultured for 24 h in the absence (-) or presence (+) of tetracycline. Proteins from 5 × 10^6^ parasites were extracted and analyzed by SDS-PAGE followed by immunoblotting against the HA epitope. Molecular mass markers are indicated in kDa in the margin.

Analysis of TbFAAH-HA localization using immunofluorescence microscopy revealed a clear perinuclear ER staining in both procyclic and bloodstream forms (Fig. 5). The signals for TbFAAH-HA surrounded the DAPI-stained nucleus and co-stained in part with the ER chaperone BiP. In bloodstream forms, TbFAAH-HA exclusively stained the perinuclear region, while in procyclic forms, weak staining was also detected in other parts of the ER (Fig. 5). The staining pattern for TbFAAH was similar to that seen for *T. brucei* ethanolamine phosphotransferase TbEPT, the enzyme catalyzing the last step in the *de novo* synthesis of ether-type PE molecular species in *T. brucei* procyclic forms (43). ER staining has also been reported for human FAAH-1 (44). In contrast, human FAAH-2 – with which TbFAAH shares more sequence homology than with FAAH-1 – has been reported to localize to lipid droplets (45).

**Figure 5:**
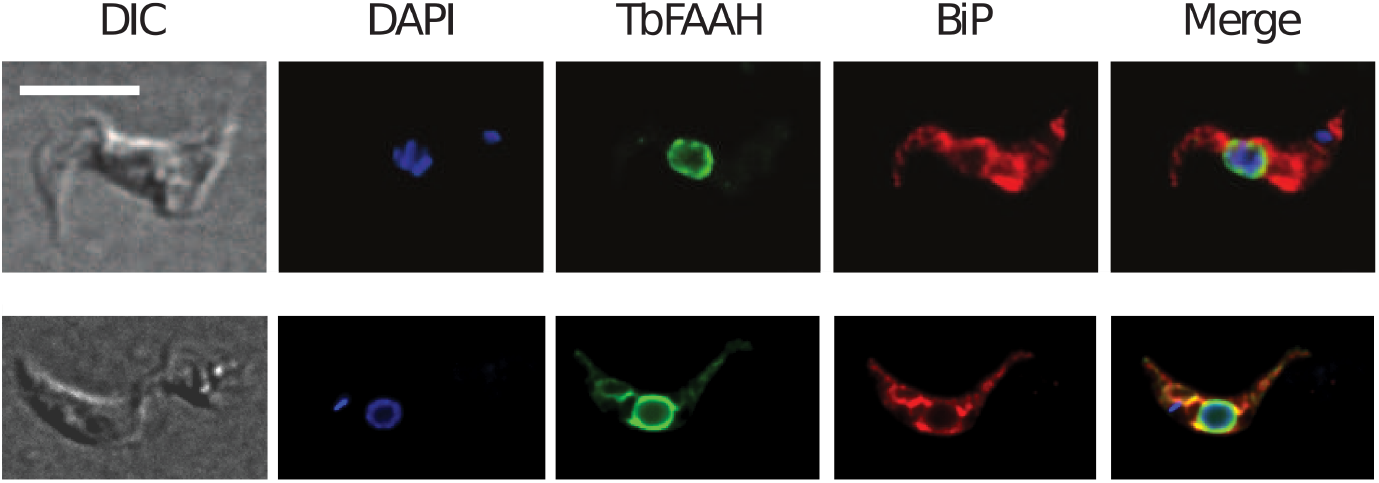
Subcellular localization of TbFAAH-HA. T. brucei bloodstream (top panels) and procyclic (bottom panels) forms expressing TbFAAH-HA were analyzed by immunofluorescence microscopy using antibodies against HA and BiP in combination with the corresponding fluorescent secondary antibodies. DNA was stained with DAPI. DIC, differential interference contrast. Scale bar: 10 µm.

### Hydrolysis of [^3^H]-AEA by *T. brucei* extracts

To study AEA hydrolysis in *T. brucei*, membrane extracts were incubated with [^3^H]-AEA and time-dependent formation of [^3^H]-ethanolamine was measured. The results show that [^3^H]-AEA was hydrolyzed by extracts from both parental (SmOx P9) and TbFAAH-KO parasites (Fig. 6), but that the activity in TbFAAH-KO parasites was consistently lower compared to parental cells. It is possible that *T. brucei* procyclic forms express more than one enzyme capable of hydrolyzing AEA. Alternatively, [^3^H]-AEA may be hydrolyzed non-specifically in the lysosome, after uptake of albumin-bound [^3^H]-AEA and trafficking to the lysosome (46).

**Figure 6:**
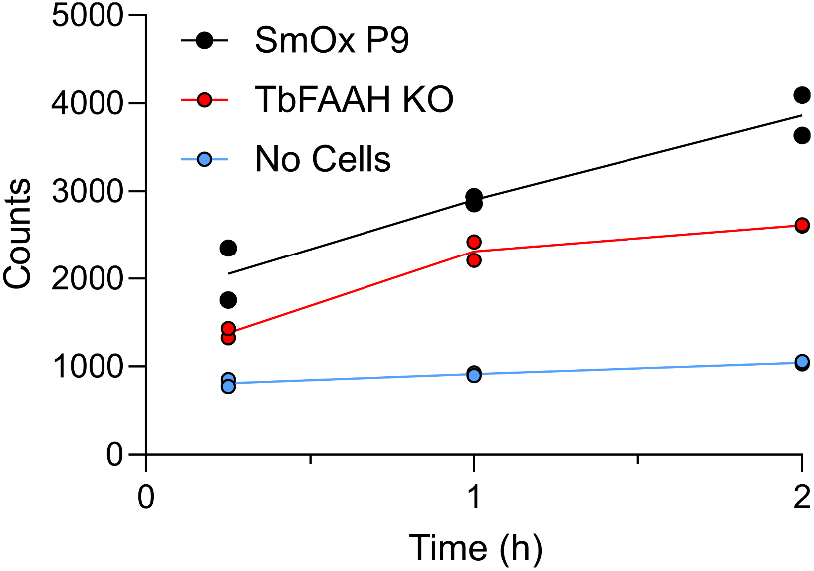
AEA hydrolysis by T. brucei membrane extracts. [^3^H]-AEA was added to membranes from equal numbers of parental (SmOx P9; black symbols) or TbFAAH-KO (red symbols) parasites and release of [^3^H]-ethanolamine was determined. Spontaneous hydrolysis of [^3^H]-AEA was determined in the absence of parasites (blue symbols).

### Incorporation of [^3^H]-AEA-derived [^3^H]-ethanolamine into PE

To study if [^3^H]-ethanolamine released from [^3^H]-AEA by TbFAAH can be used as substrate for *de novo* PE synthesis, *T. brucei* procyclic and bloodstream forms were incubated with [^3^H]-AEA and incorporation of radioactivity into PE was determined after lipid extraction by TLC and radioisotope scanning (Fig. 7). We found that both procyclic and bloodstream form parasites produced [^3^H]-PE after 24 h of incubation in the presence of [^3^H]-AEA (Fig. 7A). In addition, in line with the FAAH activity measurements, [^3^H]-PE was also formed in TbFAAH-KO cells, albeit at lower amounts than in parental parasites (Fig. 7B). No [^3^H]-ethanolamine was released from [^3^H]-AEA during 24 h of incubation in procyclic or bloodstream form culture medium in the absence of parasites. Taken together, these results demonstrate that TbFAAH-dependent hydrolysis of AEA provides ethanolamine for *de-novo* synthesis of PE in both procyclic and bloodstream form trypanosomes.

**Figure 7:**
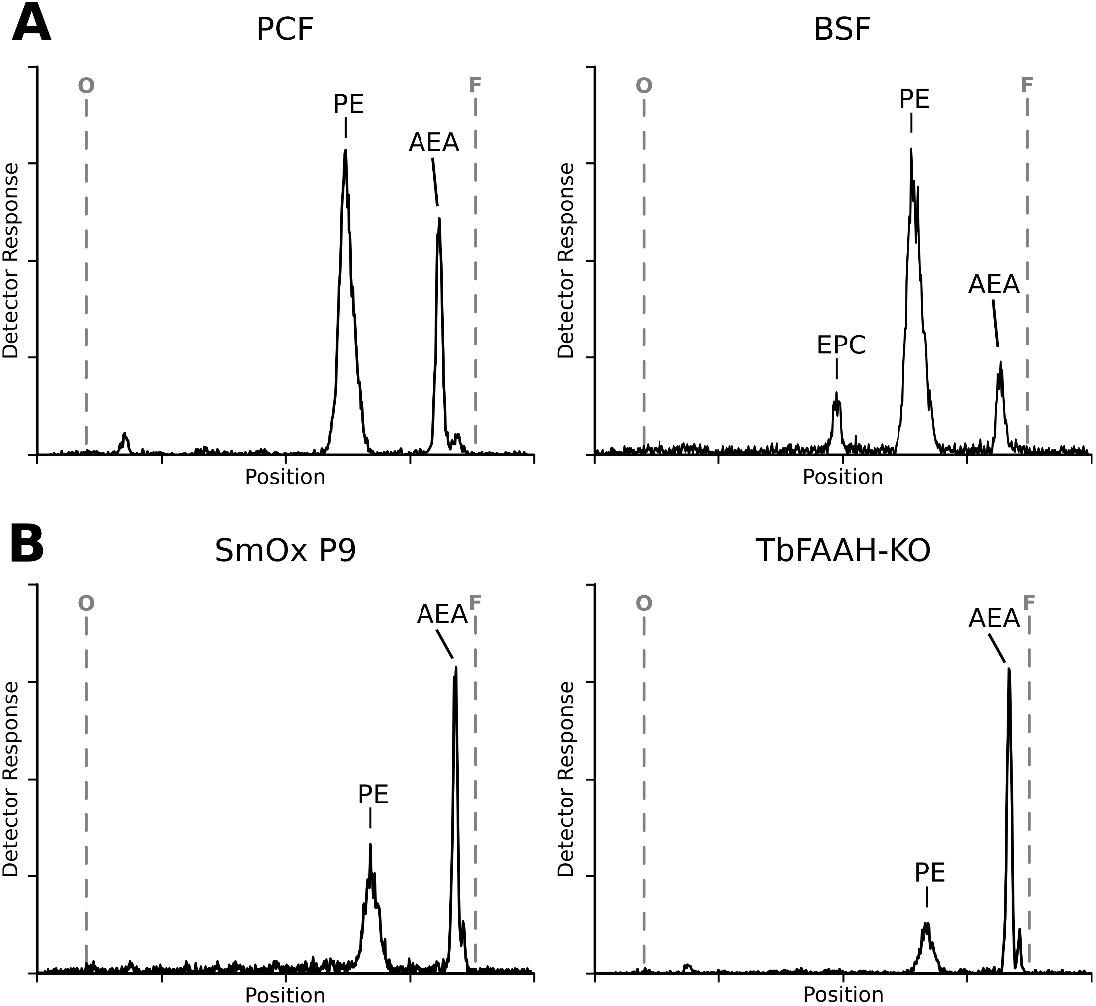
De novo synthesis of [^3^H]-PE. Equal cell numbers of T. brucei procyclic (PCF) and bloodstream (BSF) forms (A) and parental (SmOx P9) and TbFAAH-KO parasites (B) were cultured for 4 h in the presence of [^3^H]-AEA. Phospholipids were extracted and analyzed by TLC and radioisotope scanning. The positions of phosphatidylethanolamine (PE) and arachidonoylethanolamide (AEA; anandamide) were determined with radiochemically pure standards and are indicated. Ethanolamine phosphorylceramide (EPC) is only produced in bloodstream form trypanosomes (47). O, origin (site of sample application); F, solvent front.

## Acknowledgements

The work was supported by grant CRSII5_170923 from the Swiss National Science Foundation to PB. We thank Kathi Moor and Luce Farine for generating TbFAAH-HA parasites and Monika Rauch for technical assistance during parts of the study.

## Supplemental Information

**Table S1:**
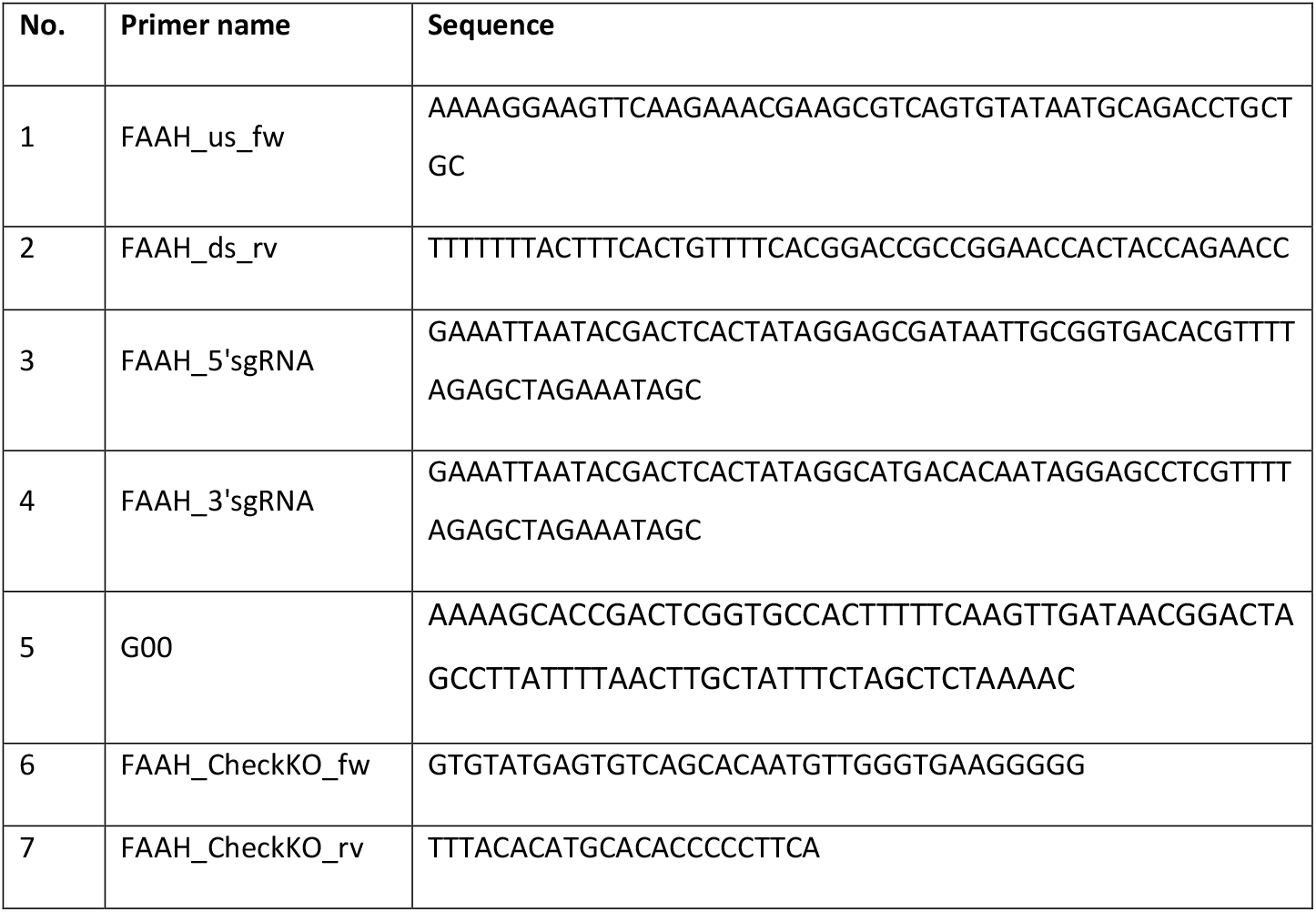
Sequences of primers used in this study.

## Bibliography

1. Vance, J. E. (2015) Phospholipid synthesis and transport in mammalian cells. Traffic. 16, 1–18

2. Bogdanov, M., Umeda, M., and Dowhan, W. (1999) Phospholipid-assisted refolding of an integral membrane protein: Minimum structural features for phosphatidylethanolamine to act as a molecular chaperone. Journal of Biological Chemistry. 274, 12339–12345

3. Tasseva, G., Bai, H. D., Davidescu, M., Haromy, A., Michelakis, E., and Vance, J. E. (2013) Phosphatidylethanolamine Deficiency in Mammalian Mitochondria Impairs Oxidative Phosphorylation and Alters Mitochondrial Morphology. Journal of Biological Chemistry. 288, 4158–4173

4. Ichimura, Y., Kirisako, T., Takao, T., Satomi, Y., Shimonishi, Y., Ishihara, N., Mizushima, N., Tanida, I., Kominami, E., Ohsumi, M., Noda, T., and Ohsumi, Y. (2000) A ubiquitinlike system mediates protein lipidation. Nature. 408, 488–492

5. Imhof, I., Canivenc-Gansel, E., Meyer, U., and Conzelmann, A. (2000) Phosphatidylethanolamine is the donor of the phosphorylethanolamine linked to the α1,4-linked mannose of yeast GPI structures. Glycobiology. 10, 1271–1275

6. Menon, A. K., Eppinger, M., Mayor, S., and Schwarz, R. T. (1993) Phosphatidylethanolamine is the donor of the terminal phosphoethanolamine group in trypanosome glycosylphosphatidylinositols. The EMBO Journal. 12, 1907

7. Signorell, A., Jelk, J., Rauch, M., and Bütikofer, P. (2008) Phosphatidylethanolamine is the precursor of the ethanolamine phosphoglycerol moiety bound to eukaryotic elongation factor 1A. Journal of Biological Chemistry. 283, 20320–20329

8. Ramakrishnan, S., Serricchio, M., Striepen, B., and Bütikofer, P. (2013) Lipid synthesis in protozoan parasites: A comparison between kinetoplastids and apicomplexans. Progress in Lipid Research. 52, 488–512

9. Smith, T. K., and Bütikofer, P. (2010) Lipid metabolism in Trypanosoma brucei. Molecular and Biochemical Parasitology. 172, 66–79

10. Kennedy, E. P., and Weiss, S. B. (1956) The function of cytidine coenzymes in the biosynthesis of phospholipides. Journal of Biological Chemistry. 222, 193–214

11. Signorell, A., Rauch, M., Jelk, J., Ferguson, M. A. J., and Bütikofer, P. (2008) Phosphatidylethanolamine in Trypanosoma brucei is organized in two separate pools and is synthesized exclusively by the Kennedy pathway. Journal of Biological Chemistry. 283, 23636–2364

12. Signorell, A., Gluenz, E., Rettig, J., Schneider, A., Shaw, M. K., Gull, K., and Bütikofer, P. (2009) Perturbation of phosphatidylethanolamine synthesis affects mitochondrial morphology and cell-cycle progression in procyclic-form Trypanosoma brucei. Mol Microbiol. 72, 1068–79

13. Gibellini, F., Hunter, W. N., and Smith, T. K. (2009) The ethanolamine branch of the Kennedy pathway is essential in the bloodstream form of Trypanosoma brucei. Molecular Microbiology. 73, 826–843

14. Samad, A., Licht, B., Stalmach, M. E., and Mellors, A. (1988) Metabolism of phospholipids and lysophospholipids by Trypanosoma brucei. Molecular and Biochemical Parasitology. 29, 159–169

15. Mellors, A., and Samad, A. (1989) The acquisition of lipids of African trypanosomes. Parasitology Today. 5, 239–244

16. Suzuki, T. T., and Kanfer, J. N. (1985) Purification and properties of an ethanolamine-serine base exchange enzyme of rat brain microsomes. Journal of Biological Chemistry. 260, 1394–9

17. Farine, L., Jelk, J., Choi, J.-Y., Voelker, D. R., Nunes, J., Smith, T. K., and Bütikofer, P. (2017) Phosphatidylserine synthase 2 and phosphatidylserine decarboxylase are essential for aminophospholipid synthesis in Trypanosoma brucei. Molecular Microbiology. 104, 412–427

18. Rifkin, M. R., Strobos, C. A. M., and Fairlamb, A. H. (1995) Specificity of Ethanolamine Transport and Its Further Metabolism in Trypanosoma brucei. Journal of Biological Chemistry. 270, 16160–16166

19. Serra, M., and Saba, J. D. (2010) Sphingosine 1-phosphate lyase, a key regulator of sphingosine 1-phosphate signaling and function. Advances in Enzyme Regulation. 50, 349–362

20. Dawoody Nejad, L., Stumpe, M., Rauch, M., Hemphill, A., Schneiter, R., Bütikofer, P., and Serricchio, M. (2020) Mitochondrial sphingosine-1-phosphate lyase is essential for phosphatidylethanolamine synthesis and survival of Trypanosoma brucei. Scientific Reports. 10, 8268

21. Elabbadi, N., Ancelin, M. L., and Vial, H. J. (1997) Phospholipid metabolism of serine in Plasmodium-infected erythrocytes involves phosphatidylserine and direct serine decarboxylation. Biochemical journal. 324 (Pt 2, 435–445

22. Farine, L., and Bütikofer, P. (2013) The ins and outs of phosphatidylethanolamine synthesis in Trypanosoma brucei. Biochim Biophys Acta. 1831, 533–542

23. Cravatt, B. F., Giang, D. K., Mayfield, S. P., Boger, D. L., Lerner, R. A., and Gilula, N. B. (1996) Molecular characterization of an enzyme that degrades neuromodulatory fattyacid amides. Nature. 384, 83–87

24. Sun, Y. X., Tsuboi, K., Zhao, L. Y., Okamoto, Y., Lambert, D. M., and Ueda, N. (2005) Involvement of N-acylethanolamine-hydrolyzing acid amidase in the degradation of anandamide and other N-acylethanolamines in macrophages. Biochim Biophys Acta. 1736, 211–220

25. Tsuboi, K., Sun, Y. X., Okamoto, Y., Araki, N., Tonai, T., and Ueda, N. (2005) Molecular characterization of N-acylethanolamine-hydrolyzing acid amidase, a novel member of the choloylglycine hydrolase family with structural and functional similarity to acid ceramidase. Journal of Biological Chemistry. 280, 11082–11092

26. Devane, W. A., Hanuš, L., Breuer, A., Pertwee, R. G., Stevenson, L. A., Griffin, G., Gibson, D., Mandelbaum, A., Etinger, A., and Mechoulam, R. (1992) Isolation and structure of a brain constituent that binds to the cannabinoid receptor. Science (1979). 258, 1946–1949

27. Lo Verme, J., Fu, J., Astarita, G., La Rana, G., Russo, R., Calignano, A., and Piomelli, D. (2005) The nuclear receptor peroxisome proliferator-activated receptor-α mediates the anti-inflammatory actions of palmitoylethanolamide. Molecular Pharmacology. 67, 15–19

28. Gachet, M. S., Rhyn, P., Bosch, O. G., Quednow, B. B., and Gertsch, J. (2015) A quantitiative LC-MS/MS method for the measurement of arachidonic acid, prostanoids, endocannabinoids, N-acylethanolamines and steroids in human plasma. Journal of Chromatography B: Analytical Technologies in the Biomedical and Life Sciences. 976–977, 6–18

29. Marazzi, J., Kleyer, J., Paredes, J. M. V., and Gertsch, J. (2011) Endocannabinoid content in fetal bovine sera - Unexpected effects on mononuclear cells and osteoclastogenesis. Journal of Immunological Methods. 373, 219–228

30. Beneke, T., Madden, R., Makin, L., Valli, J., Sunter, J. D., and Gluenz, E. (2017) A CRISPR Cas9 high-throughput genome editing toolkit for kinetoplastids. Royal Society Open Science. 4, 170095

31. Wirtz, E., Leal, S., Ochatt, C., and Cross, G. A. M. (1999) A tightly regulated inducible expression system for conditional gene knock-outs and dominant-negative genetics in Trypanosoma brucei. Molecular and Biochemical Parasitology. 99, 89–101

32. Dean, S., Sunter, J. D., Wheeler, R. J., Hodkinson, I., Gluenz, E., and Gull, K. (2015) A toolkit enabling efficient, scalable and reproducible gene tagging in trypanosomatids. Open Biology. 5, 140197

33. Serricchio, M., and Bütikofer, P. (2013) Phosphatidylglycerophosphate synthase associates with a mitochondrial inner membrane complex and is essential for growth of Trypanosoma brucei. Molecular Microbiology. 87, 569–579

34. Nicolussi, S., Viveros-Paredes, J. M., Gachet, M. S., Rau, M., Flores-Soto, M. E., Blunder, M., and Gertsch, J. (2014) Guineensine is a novel inhibitor of endocannabinoid uptake showing cannabimimetic behavioral effects in BALB/c mice. Pharmacological Research. 80, 52–65

35. Bligh, E. G., and Dyer, W. J. (1959) a Rapid Method of Total Lipid Extraction and Purification. Canadian Journal of Biochemistry and Physiology. 37, 911–917

36. Chicca, A., Marazzi, J., Nicolussi, S., and Gertsch, J. (2012) Evidence for bidirectional endocannabinoid transport across cell membranes. Journal of Biological Chemistry. 287, 34660–82

37. Wei, B. Q., Mikkelsen, T. S., McKinney, M. K., Lander, E. S., and Cravatt, B. F. (2006) A second fatty acid amide hydrolase with variable distribution among placental mammals. Journal of Biological Chemistry. 281, 36569–36578

38. Chebrou, H., Bigey, F., Arnaud, A., and Galzy, P. (1996) Study of the amidase signature group. Biochim Biophys Acta. 1298, 285–93

39. Dean, S., Sunter, J. D., and Wheeler, R. J. (2017) TrypTag.org: A Trypanosome Genome-wide Protein Localisation Resource. Trends Parasitol. 33, 80–82

40. Liu, Y., Patricelli, M. P., and Cravatt, B. F. (1999) Activity-based protein profiling: the serine hydrolases. Proc Natl Acad Sci U S A. 96, 14694–9

41. Patricelli, M. P., Giang, D. K., Stamp, L. M., and Burbaum, J. J. (2001) Direct visualization of serine hydrolase activities in complex proteomes using fluorescent active sitedirected probes. Proteomics. 1, 1067–71

42. Mor, M., Rivara, S., Lodola, A., Plazzi, P. V., Tarzia, G., Duranti, A., Tontini, A., Piersanti, G., Kathuria, S., and Piomelli, D. (2004) Cyclohexylcarbamic acid 3’-or 4’-substituted biphenyl-3-yl esters as fatty acid amide hydrolase inhibitors: Synthesis, quantitative structure-activity relationships, and molecular modeling studies. Journal of Medicinal Chemistry. 47, 4998–5008

43. Farine, L., Niemann, M., Schneider, A., and Bütikofer, P. (2015) Phosphatidylethanolamine and phosphatidylcholine biosynthesis by the Kennedy pathway occurs at different sites in Trypanosoma brucei. Scientific Reports. 5, 16787

44. Kaczocha, M., Glaser, S. T., and Deutsch, D. G. (2009) Identification of intracellular carriers for the endocannabinoid anandamide. Proceedings of the National Academy of Sciences. 106, 6375–80

45. Kaczocha, M., Glaser, S. T., Chae, J., Brown, D. A., and Deutsch, D. G. (2010) Lipid droplets are novel sites of N-acylethanolamine inactivation by fatty acid amide hydrolase-2. Journal of Biological Chemistry. 285, 2796–2806

46. Bojesen, I. N., and Hansen, H. S. (2003) Binding of anandamide to bovine serum albumin. J Lipid Res. 44, 1790–4

47. Sutterwala, S. S., Hsu, F. F., Sevova, E. S., Schwartz, K. J., Zhang, K., Key, P., Turk, J., Beverley, S. M., and Bangs, J. D. (2008) Developmentally regulated sphingolipid synthesis in African trypanosomes. Molecular Microbiology. 70, 281–296

